# Controlled spatial organization of bacterial clusters reveals cell filamentation is vital for *Xylella fastidiosa* biofilm formation

**DOI:** 10.1101/2021.01.08.425936

**Authors:** Silambarasan Anbumani, Aldeliane M. da Silva, Eduarda Regina Fischer, Mariana de Souza e Silva, Antônio A.G. von Zuben, Hernandes F. Carvalho, Alessandra A. de Souza, Richard Janissen, Monica A. Cotta

## Abstract

The morphological plasticity of bacteria to form filamentous cells commonly represents an adaptive strategy induced by stresses. In contrast, for diverse pathogens filamentous cells have been observed during biofilm formation, with function yet to be elucidated. To identify prior hypothesized quorum sensing as trigger of such cell morphogenesis, spatially controlled cell adhesion is pivotal. Here, we demonstrate highly-selective cell adhesion of the biofilm-forming phytopathogen *Xylella fastidiosa* to gold-patterned SiO_2_ substrates with well-defined geometries and dimensions. The consequent control of both cell density and distances between cell clusters using these patterns provided evidence of quorum sensing governing filamentous cell formation. While cell morphogenesis is induced by cell cluster density, filamentous cell growth is oriented towards neighboring cell clusters and distance-dependent; large interconnected cell clusters create the early biofilm structural framework. Together, our findings and investigative platform could facilitate therapeutic developments targeting biofilm formation mechanisms of *X. fastidiosa* and other pathogens.

## INTRODUCTION

Bacteria have evolved diverse surface adhesion mechanisms to enable biofilm formation on biotic and abiotic substrates in a variety of natural, medical, and industrial settings^1–3^. The virulence of pathogenic bacteria strongly depends on their capability to attach to biotic surfaces and form multicellular assemblies^4–8^. Such pathogenic biofilms are highly resistant to diverse antimicrobial compounds due to their encapsulation within a matrix of hydrated extracellular polymeric substances (EPS)^7,9^. A better understanding of the mechanisms of bacterial adhesion and biofilm formation is thus vital to reveal potential vulnerabilities that can lead to their prevention and disruption^2,10^. In our previous work we characterized the individual stages in the process of biofilm formation in *Xylella fastidiosa*^11^, a vascular phytopathogen that causes large economical damage worldwide by inducing diseases in a range of important crops (e.g. citrus, grape, coffee, almond, olives, among others)^12,13^ and further shares genetic traits with human biofilm-forming pathogens^14,15^. With respect to the vital question of how large-sized biofilms are formed by this pathogen, we previously observed that cells during biofilm growth elongated up to 10-fold their typical size when connected with neighboring bacterial clusters, which we describe as a macroscale biofilm framework^11^. In this scenario, the extreme elongation of cells represents a central feature of biofilm formation, rather than simply a consequence of stresses such as starvation and DNA damage as commonly observed in other bacteria^16–19^. While similar filamentous cells have also been observed in *Vibrio cholera*^6^, *Caulobacter crescentus*^18^ and *Pseudomonas aeruginosa*^20^ during biofilm formation, solid evidence that such elongated cells are necessary for triggering or development of biofilms remains fragmentary.

Whether filamentous cell growth of *X. fastidiosa* is the trigger rather than a consequence of biofilm formation remains a central question. Earlier results revealed that only a small fraction of cells undergo morphogenesis to filamentous cells^11^, which emanate from bacterial clusters. In this scenario, a stress-based trigger for filamentation does not seem to be present since such a stress would likely be shared by most cells in the cluster, and would therefore promote the morphogenesis of a majority of cells^16,17^. Another potential mechanism of triggering morphogenesis poses cell-cell communication *via* diffusible signaling factors (DSF) as part of the quorum sensing system. In particular, *X. fastidiosa* regulates its cell-cell adhesion, biofilm formation, and virulence in a cell density-dependent manner *via* DSF-based signaling^21–24^. A prime example of such quorum sensing regulation, the expression of adhesins as well as the secretion of outer membrane vesicles (OMVs), which contain biologically active biomolecules associated with cell functions linked to cell adhesion and virulence, occurs in a DSF-dependent, apparently cell density-dependent fashion^25–27^. Both adhesin expression and OMVs secretion in *X. fastidiosa* modulate its systematic dissemination in the host^27,28^. Therefore, since filamentous cells have been associated with biofilm formation, which itself is regulated by cell-cell communication^11^, we hypothesize that their formation is also a DSF-dependent process.

Since signaling molecules can strongly affect bacterial physiology and virulence, several methods for their *in vitro* and *in vivo* detection have been previously developed^29,30^. The majority of these methods rely on chromatographic and mass spectrometric techniques, or biosensor systems using genetically modified reporter bacteria^29^. However, such approaches include important limitations, particularly for signaling molecules used by *X. fastidiosa*. The detection of DSF molecules secreted by a small number of cells, such as when clusters are comprised of only a few cells, is difficult and requires methods capable of reliably detecting very low DSF concentrations. As such, assessing spatial gradients associated with filamentous cell formation poses an extreme challenge.

In this work, we circumvented these bottlenecks and elucidated the triggering mechanisms for *X. fastidiosa* cell filamentation during biofilm formation. Since the local DSF concentration produced by bacterial clusters in a non-continuous liquid flow system decreases gradually and isotropically with distance, controlled spatial separation of bacterial adhesion is critical to ascertain its role in any process involving quorum sensing. Our previous works have suggested that *X. fastidiosa* has a higher adhesion affinity to gold than to other abiotic and biotic chemical substrates^4,11,31^. Here, we confirmed the high adhesion affinity of *X. fastidiosa* to gold and demonstrated that its spatial organization can be controlled by using lithographically-defined gold patterns, therefore enabling the modulation of the size and distance between bacterial clusters. Exploiting its strong adhesion on gold, we were able to probe the formation of filamentous cells over an 18-hour period and determine the effect of cell aggregate sizes and distance between spatially separated bacterial clusters. We demonstrated that the formation of filamentous cells is induced by local bacterial density, and that their growth is oriented toward neighboring cell clusters in a distance-dependent manner, which eventually creates a network of interconnected cell clusters. Our results provide not only evidence that filamentous cell formation depends on cell density, but also that the resulting formation of large-size biofilm frameworks is governed by a quorum sensing process, which may lead to novel translational schemes to inhibit biofilm-forming pathogens.

## RESULTS

### *X. fastidiosa* preferentially adheres to gold surfaces

Since our previous work^4,11^ indicated that *X. fastidiosa* adheres more efficiently to gold than to diverse other biotic and abiotic surfaces, we first compared the propensity *of X. fastidiosa* to adhere to Au rather than to SiO_2_ as a function of time using quantitative assays. We fabricated Au micro patterns on SiO_2_ surfaces with different shapes and dimensions using Direct-Write Laser (DWL) photolithography, followed by deposition of a 20 nm thick Au coating using e-beam evaporation (**Figure 1**). After photo resist lift-off and cleaning, the substrates were sterilized with oxygen plasma prior to bacterial growth experiments.

**Figure 1.**
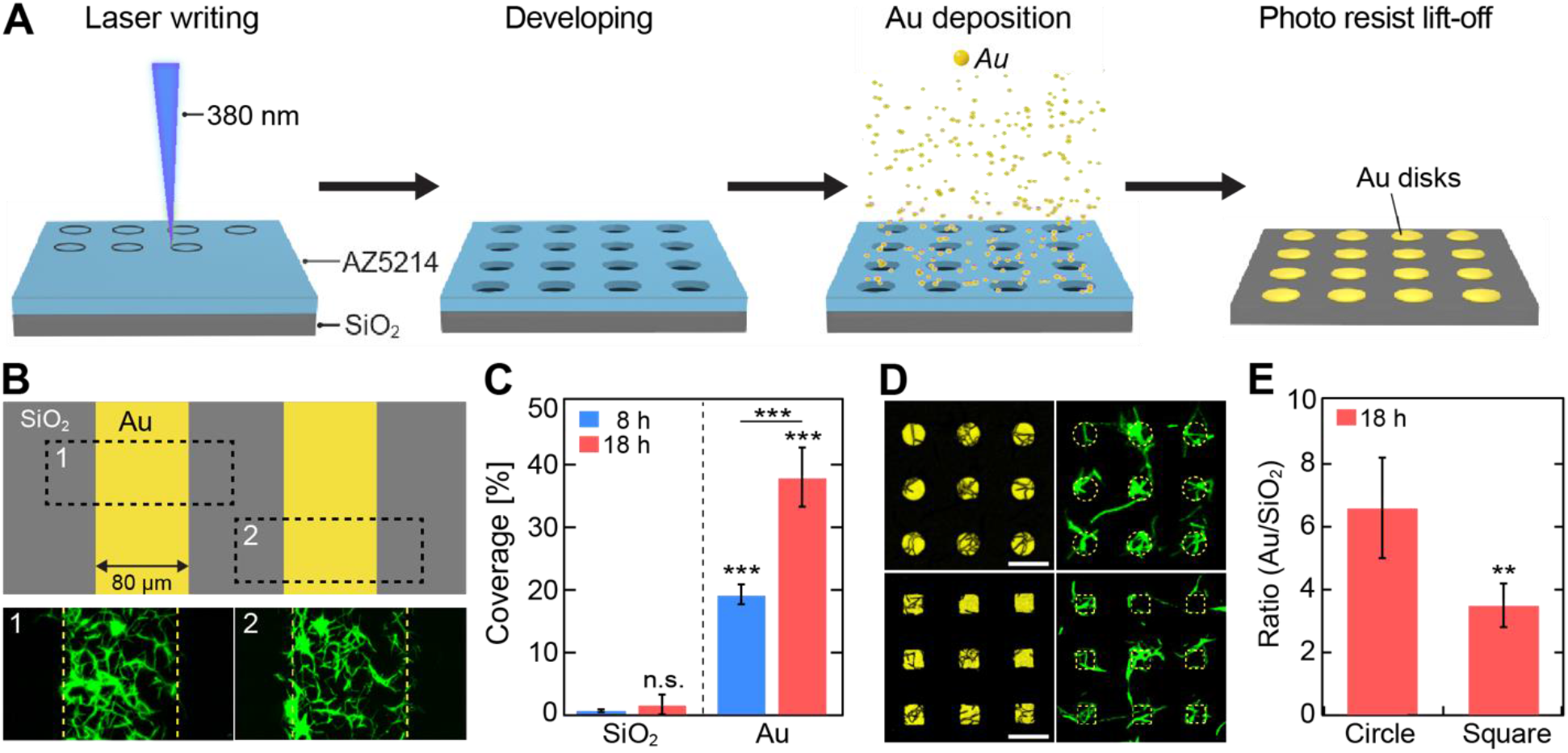
Fabrication of Au micro patterns and substrate-selective bacterial adhesion. (**A**) Stepwise schematic representation of Au pattern fabrication on SiO_2_ substrate. (**B**) Representative fluorescence images of selective bacterial adhesion to Au line patterns and SiO_2_ substrate after 18 h growth. (**C**) Bacteria coverage on SiO_2_ and Au line patterns after 8 h and 18 h of growth. (**D**) Representative fluorescence and corresponding laser reflection images of bacterial adhesion on circular- (11 µm diameter) and square-shaped (11 µm edge length) Au arrays, separated by 9 µm, after 18 h growth; scale bar denotes 20 µm. (**E**) Ratio of Au to SiO_2_ bacterial coverage on circular and square shaped Au arrays after 18 h growth. Asterisks indicate statistical significance (**p < 0.01; ***p < 0.001; n.s. = non-significant) resulting from two-tailed, unpaired t-tests.

On large Au areas with dimensions of 80 µm x 12 mm (**Figure 1B**, top), we incubated GFP-expressing *X. fastidiosa* strain 1139911,32 for 8 and 18 h, after which non-attached bacteria were removed by gentle rinsing. Strikingly, widefield fluorescence microscopy (WFM) images revealed (**Figure 1B**, bottom) that cells predominantly adhered to Au surfaces, even after 18 h of growth. The quantification of the bacterial coverage of equal areas of SiO_2_ and Au (**Figure 1C**) revealed that significantly more (10-20-fold) cells adhered to Au as compared to SiO_2_, independent of growth duration. While cell adhesion to SiO_2_ was comparably low (< 2% surface coverage) at both growth durations, a 2-fold higher bacterial coverage was observed after 18 h growth on Au compared to 8 h. These results not only support the previous finding that *X. fastidiosa* predominantly adheres to Au, but also provide the means to create spatial patterns of bacterial colonies with controlled spatial separation.

Circular- and square-shaped Au micro patterns were created (**Figure 1D**, left) to probe whether particular geometries facilitate cell adhesion. These patterns were incubated with *X. fastidiosa* for an extended time of 18 h, since the growth duration did not affect the preferential adhesion of *X. fastidiosa* to Au (**Figure 1C**); furthermore, an increased number of attached cells after extended growth duration time would facilitate their detection. Confocal Laser Scanning Microscopy (CLSM) of these samples (**Figure 1D**, right) also showed predominant cell adhesion to Au-coated regions. Importantly, the formation of elongated, filamentous cells was readily observable on both Au pattern geometries (**Figure 1D**).

In order to quantitatively determine the bacterial coverage on both Au pattern geometries, either CLSM reflective images or WFM bright field images, together with their corresponding fluorescence microscopy images, were processed into binary images (**Supplementary Figure S1A-D**) and subtracted from each other to determine the cell coverage on Au and SiO_2_ separately. To compare shape-dependent cell adhesion propensities, we calculated the cell coverage ratios of Au to SiO_2_ (**Figure 1E**) from the cell coverages measured for each shape pattern (**Supplementary Figure S1E**). The results (**Figure 1E**) clearly show a significantly higher cell coverage ratio for circular-shaped than for square-shaped Au arrays. Circular-shaped Au arrays were thus used to probe distance- and density-dependent formation of filamentous cells.

### Spatial proximity and density of bacterial clusters induce cell elongation

The effect of growth duration, Au disk diameter and distance on both the cell adhesion efficiency and substrate-dependent cell adhesion specificity was assessed. We probed samples grown for longer than that of the typical *X. fastidiosa* division time^11^ of ∼ 6 h, namely 6, 8, 14 and 18 h. Using an Au disk diameter of 11 µm, as used in our substrate-selectivity experiments (**Figure 1D** and **1E**), we simultaneously examined the effect of different growth durations and Au pattern distances. Here, we used 9 and 14 µm separation distances, representing values larger than approx. 2-fold and 3-fold the typical length of *X. fastidiosa* cells (∼3-4 µm^12^), respectively. We reasoned by this approach to be able to readily discriminate between normal cell lengths and those of filamentous cells. Remarkably, independent of growth durations and Au pattern distances, the propensity of cells to adhere to Au rather than SiO_2_ remained (**Figure 2A**).

**Figure 2:**
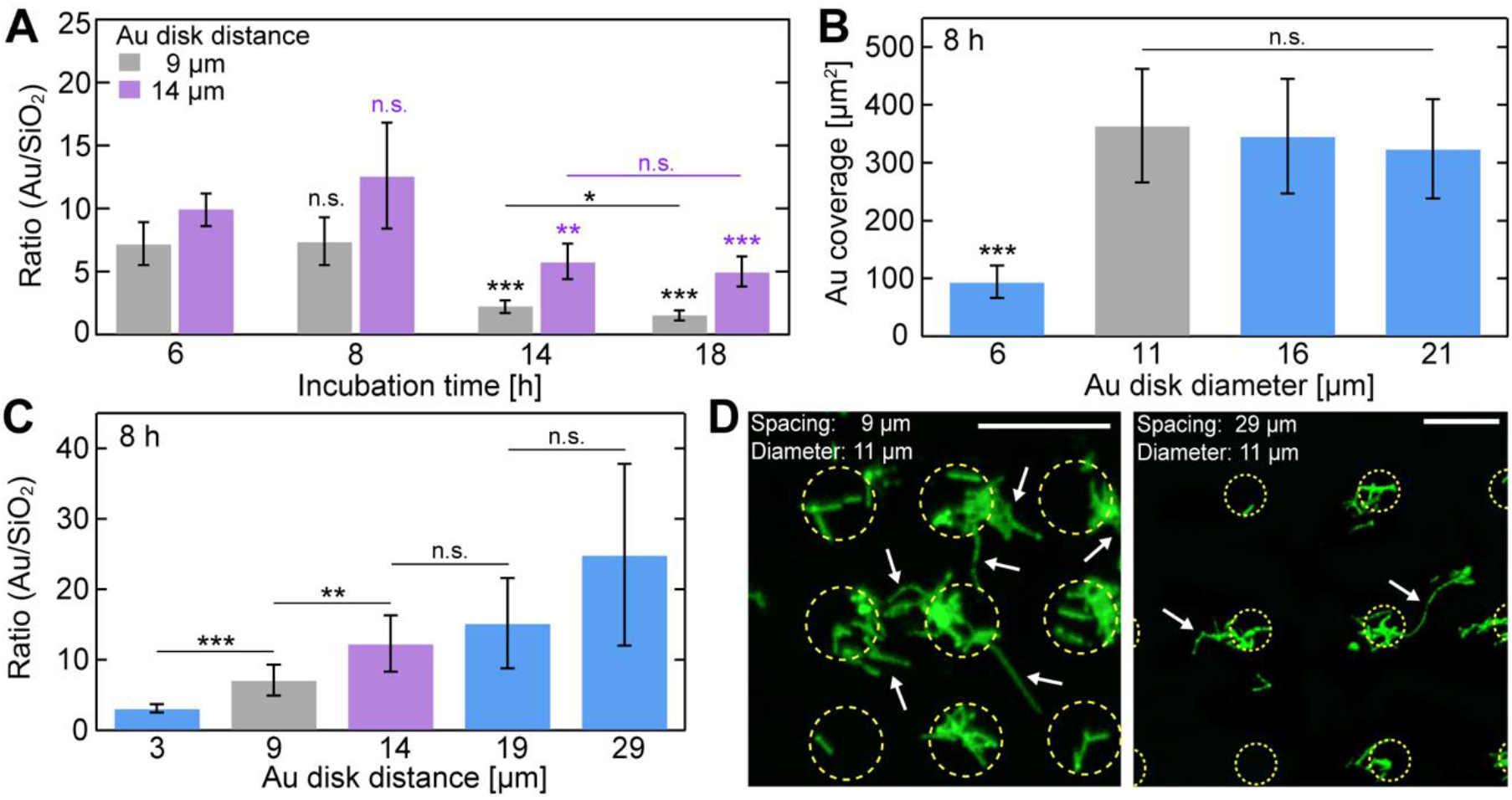
Substrate-selectivity of bacterial adhesion in dependency of growth time, Au disk diameter, and Au pattern distance. (**A**) Bacterial adhesion ratio of Au to SiO_2_ of circular Au disks arrays with 11 µm diameter, separated by 9 and 14 µm spacing for different growth times (6, 8, 14, 18 h). (**B**) Bacterial coverage on different Au disk diameters (6, 11, 16, 21 µm) with 9 µm spacing for 8 h growth time. (**C**) Bacterial adhesion ratio of Au to SiO_2_ of circular Au disks arrays with 11 µm diameter, separated by different spacing (3, 9, 14, 19, 29 µm) for 8 h growth time. (**D**) Representative fluorescence images of *X. fastidiosa* cells adhering to circular Au disks arrays with 11 µm diameter,separated by 9 µm (left) and 29 µm (right) spacing for 8 h growth time; scale depicts 20 µm. Asterisks indicate statistical significance (*p < 0.05; **p < 0.01; ***p < 0.001; n.s. = non-significant), resulting from two-tailed, unpaired t-tests.

However, when the typical *X. fastidiosa* division time of ∼6 h was largely exceeded (14 and 18 h incubation), a considerable number of ≥3^rd^ generation daughter cells were encountered on the SiO_2_ substrate, and the substrate selectivity seemed reduced. Since the cell adhesion ratios of Au relative to SiO_2_ between 6 and 8 h, and between 14 and 18 h, were statistically comparable for both Au pattern distances, subsequent experiments were carried out using 6-8 h and 18 h growth durations. We reasoned that the 6-8 h growth duration may resemble the initial stages of biofilm formation dictated primarily by adhesion of planktonic cells, whereas 18 h reflected the detection of slow-growing filamentous cells as the cell mass adhering to SiO_2_ (e.g. **Figure 1D**).

Yet another possibility causing *X. fastidiosa* cells adhering to SiO_2_ could result from a limiting Au area for further planktonic and daughter cell adhesion, resulting in increased adherence to SiO_2_. To verify this possibility, we varied the Au disk diameter in our arrays, ranging from 6 µm to 21 µm in increasing steps of 5 µm (which corresponds to the upper limit for typical cell lengths^12^). Indeed, while Au disks with diameters ≥11 µm exhibited similar cell coverage (∼370 µm^2^), smaller disks with a diameter of 6 µm (**Figure 2B**) showed a 4-fold lower cell coverage (∼90 µm^2^). Although expected from the different disk areas, this result also indicated that small Au disk areas can limit bacterial coverage; we proceeded with 11 µm disk diameters for all subsequent experiments, since this limiting effect was not observable for Au disk diameters ≥11 µm.

If indeed potential cell-cell communication *via* quorum sensing mediates the formation of filamentous cells^7,33,34^, we reasoned that variances in distance between Au disks should affect this process. Following this idea, the formation of filamentous cells was monitored *via* fluorescence imaging of cells clusters separated over various distances. Therefore, *X. fastidiosa* was incubated on Au disk patterns with distances ranging from 3 to 29 µm (**Figure 2C**). As observed before, the cell adhesion propensity was higher on Au than on the SiO_2_ substrate, but, in turn, the cell coverage ratio of Au to SiO_2_ also increased with separation distance. This result indicates that the cell mass adhering to SiO_2_ between the Au disks, which predominantly consists of both elongated and filamentous cells (**Figure 2D**), decreased with distance. A distance of 3 µm between Au disks was unusable in the study of the formation of filamentous cells since the typical cell size of 3-4 µm^12^ already bridged the SiO_2_ area between neighboring Au disks upon cell adherence (**Supplementary Figure S2**). More strikingly, significant differences in the Au to SiO_2_ ratios were encountered between 9 and 14 µm disk separation, but not in the case of larger distances (19 and 29 µm). This observation rendered Au disk distances of 9 and 14 µm ideal for systematic investigation of bacterial cluster proximity and size in the formation of filamentous cells.

### Formation of filamentous cells depends on bacterial cluster density and distance

To evaluate whether the density of cell aggregates or their proximity is the decisive parameter for filamentous cell formation, we analyzed all *X. fastidiosa* cell lengths in samples with 9 and 14 µm Au disk distances, and grown over 6 and 18 h. Notably, the cell length distribution over all samples (**Figure S3A**) revealed the existence of three distinct cell length populations. The cell length distributions for each of the four tested conditions (**Figure 3A**) exhibited a similar picture with three populations of comparable average length values. After 6 h of cell growth (**Figure 3A**, top graphs), where predominantly planktonic cells adhere and at most one cell division occurs, the dominant population exhibited an average cell length of ca. 3-4 µm, independent of the Au disk distance (**Figure 3B**, left).

**Figure 3:**
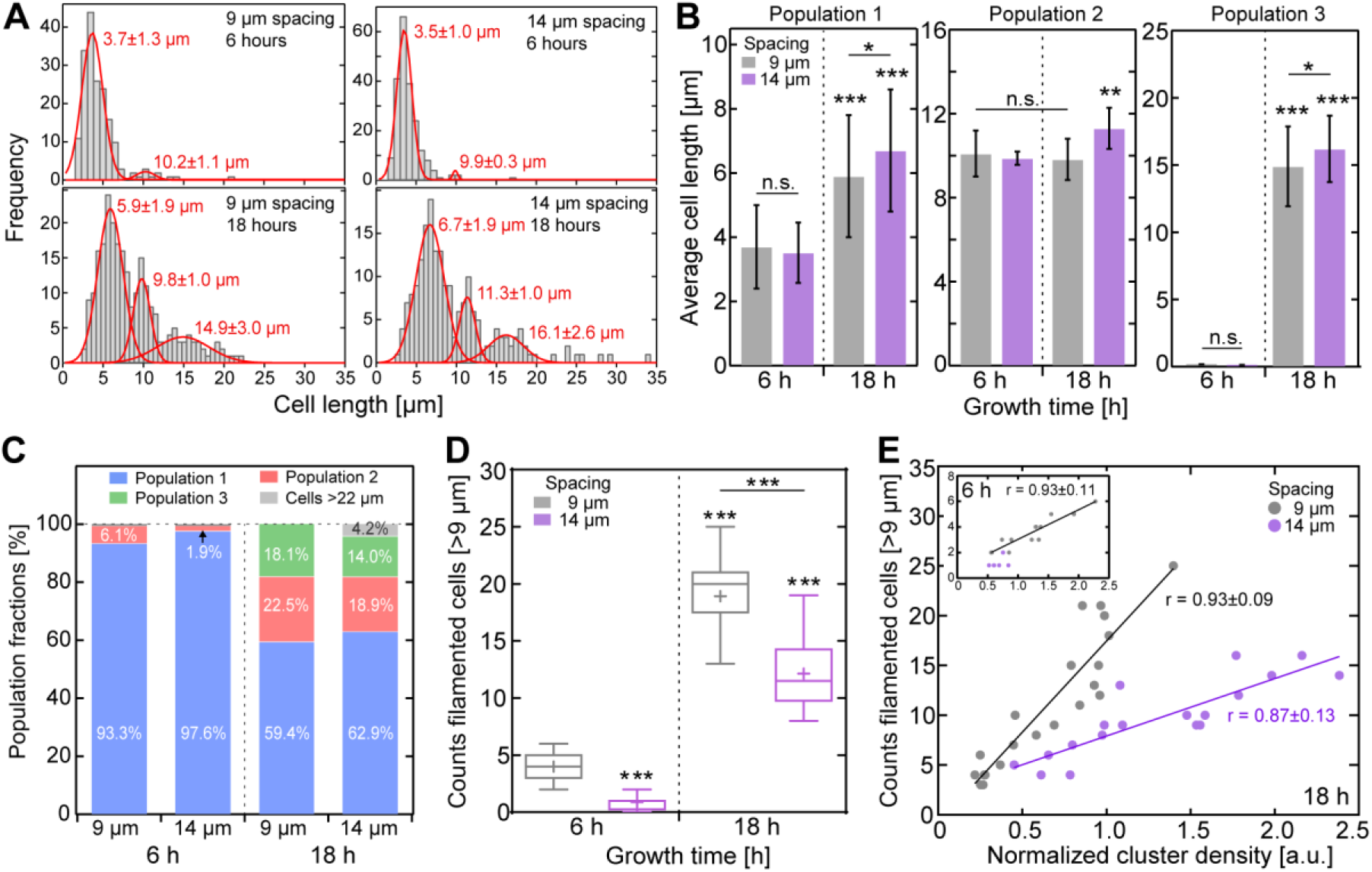
Bacterial cluster distance- and density-dependent formation of filamentous cells. (**A**) Cell length distributions for 9 and 14 µm spacing, grown over 6 and 18 h (6 h: 9 µm, *N* = 183; 14 µm, *N* = 213; 18h: 9 µm, *N* = 249; 14 µm, *N* = 191). Gaussian fits (red) resulting from the applied Gaussian mixture exhibit three distinct cell length populations (see also Supplementary Figure S3A); the fit values (±SD) are denoted within the plot. (**B**) Comparison of average cell lengths for each population (AVG±SD). (**C**) Fractions (integral of Gaussian fits) of the different populations observed in all tested conditions. (**D**) Number of filamentous cells detected in the proximity of - or emanating from - bacterial clusters for all tested conditions (from *N* = 9 fluorescence images each). (**E**) Number of filamentous cells in dependency of cluster density after 18 h growth (inset for 6h growth), calculated from the integral of bacterial cluster fluorescence intensity and normalized to single filamentous cells for all tested conditions. Lines represent linear fits and the corresponding Pearson correlation coefficients *r* are denoted for each fit. Asterisks indicate statistical significance (***p < 0.001), resulting from two-tailed, unpaired t-tests.

While this average cell length is comparable to that of a typical *X. fastidiosa* cell^12^, a minor secondary population of filamentous cells were found with an average cell length of ∼10 µm (**Figure 3B**, center). The increase in cell length was more distinct after 18 h cell growth (**Figure 3A**, bottom): while the subpopulation with average cell lengths ∼10 µm strongly increased in proportion, a third population of filamentous cells with an average cell length of ∼15-16 µm appears (**Figure 3C**). This third subpopulation exhibited a 4-fold increase in cell length compared to a typical *X. fastidiosa* cell of 3-4µm (**Figure 3B**, right).

Interestingly, the average cell length after 18 h growth exhibited strong distance-dependent increase, with significantly larger cells observed for samples separated by 14 µm (**Figure 3B**). Under this condition, an additional small subpopulation of very long filamentous cells with lengths>22 µm emerged (**Figure 3A**, bottom right; **Figure 3C**). Given that 14 µm pattern distance results in a diagonal distance between the Au disks of ∼20 µm, the appearance of filamentous cells of such lengths was to be expected. Both the increase in filamentous cell length and the appearance of very long cells >20 µm in case of 14 µm pattern distance might imply that the growth of filamentous cells, predominantly emanating from cluster boundaries, might be directed towards and connect adjacent clusters, as we indeed observed in fluorescence images (**Supplementary Figure S3B**).

We next sought out to determine whether the formation of filamentous cells depends on growth duration and spatial distance between cell clusters. For this analysis, it was necessary to define a cell length that allows to discriminate cells as being filamentous or not. Since the dominant cell length population after only 6 h of cell growth predominantly represents non-elongated cells with sizes typically encountered for *X. fastidiosa*^12^, the second population, which increases in relative abundance after extended growth duration, was deemed to consist largely of filamentous cells. We thus defined cells with lengths of 9 µm or larger, corresponding to the one-sigma lower bound of the second population distribution (**Supplementary Figure S3A**), as filamentous cells. Upon quantifying the number of filamentous cells in all tested conditions, we found up to a 5-fold higher abundance of filamentous cells after 18 h growth than we observed after 6 h growth (**Figure 3D**). Importantly, while longer growth duration increased the occurrence of filamentous cells independent from spatial separation, the number of filamentous cells notably decreased with spatial distance between cell clusters.

Since extended growth duration led to an increase of filamentous cell formation, we questioned whether the cell density that increases with time, rather than growth duration itself, is the decisive parameter. To address this question, we estimated the bacterial cluster densities by integrating their fluorescence intensity by taking into account that integrated fluorescence intensity scales linearly with cell density. This method was also deemed appropriate since the long *X. fastidiosa* cell-division time of ∼6 h^11^ allows the observation of the horizontal expansion of cell within our observation times before the formation of large 3D biofilm architectures. Interestingly, our results revealed (**Figure 3E**) a strong correlation (Pearson correlation coefficient *r* ∼0.9) between the abundance of filamentous cells and the abundance of cells within clusters. However, and more importantly, while there was no effect of growth duration on filamentous cell formation observable, we detected that the distance between clusters had a significant effect (**Figure 3E)** on filamentous cell formation. Larger distances between clusters (14 µm *vs* 9 µm) resulted in significantly less filamentous cells, even at high cell densities. These results confirm our concept that filamentous cell formation depends on both cell density and their spatial separation.

### Filamentous cells interconnect bacterial clusters to form biofilm frameworks

Our previous work had raised the concept that the underlying function of filamentous cells is to connect spatially separated cell aggregates to form the macroscale network that is required for the formation of subsequent large-scale mature biofilms^11^. Our observation of filamentous cells that interconnect adjacent bacterial clusters (**Figure 4A**) supports this model. We reasoned that the growth of filamentous cells might be directed towards adjacent bacterial clusters and is potentially governed by a quorum sensing mechanism, triggered by locally high concentrations of DSF in the surrounding environment of clusters. If this model is true, we would expect that both cell cluster size and the distance between clusters would drive filamentous cells to connect adjacent cell aggregates. To verify this model, we differentiated between *formed* filamentous cells (originating from single cell clusters) and *interconnecting* (bridging two or more cell clusters) filamentous cells. Since we found that filamentous cell formation depends on cell cluster size and separation distance, we determined the occurrence of the two classes of filamentous cells as a function of these two parameters.

**Figure 4.**
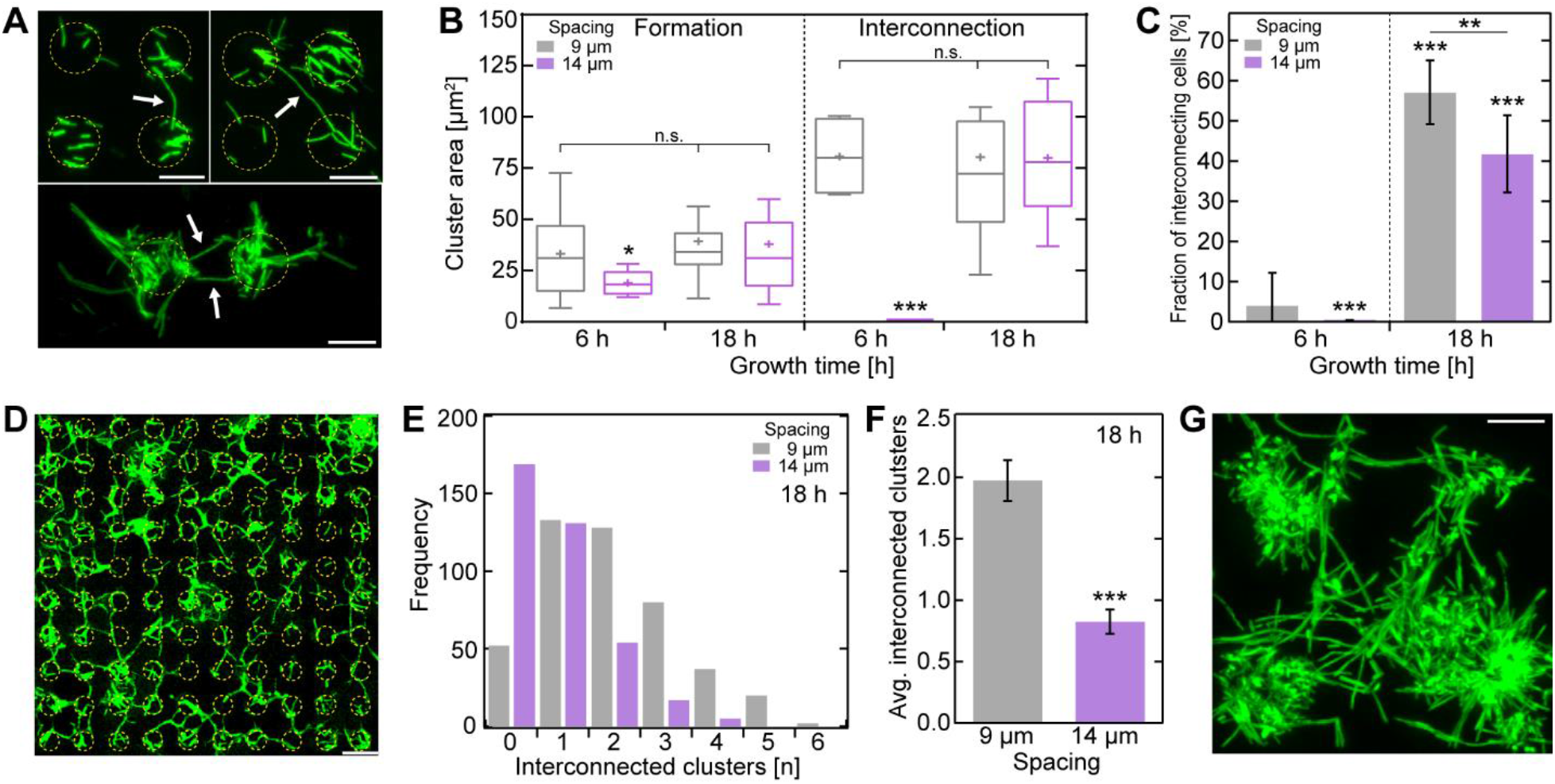
Filamentous cell growth is directed to adjacent bacterial clusters and interconnect cell clusters to a large network. (**A**) Example fluorescence images of bacterial cluster pairs interconnected by filamentous cells. Filamentous cells are indicated by arrows; scale bar depicts 10 µm. (**B**) Bacterial cluster area-dependent *formation* of filamentous cells and filamentous cells that *interconnect* adjacent bacterial clusters for 9 and 14 µm spacing samples, grown for 6 and 18 h. The cluster size is normalized to one filamentous cell (if multiple). (**C**) Fraction of *interconnecting* cells of all filamentous cells observed for 9 and 14 µm distances, grown for 6 and 18 h. (**D**) Example fluorescence image of an Au disk array with 9 µm spacing after 18 h growth shows multiple interconnected bacterial clusters; scale bar depicts 20 µm. (**E**) Degree of interconnected clusters (*N* = 446 for 9 µm, *N* = 372 for 14 µm distances) and (**F**) average number (± 95% confidence interval) of cluster interconnections of 9 and 14 µm distances arrays after 18 h growth. (**G**) I*n-vitro* CLSM fluorescence image of growing biofilm, originating from clusters interconnected with multiple filamentous cells; scale bar depicts 20 µm. Asterisks indicate statistical significance (*p < 0.05; **p < 0.01; ***p < 0.001; n.s. = non-significant), resulting from two-tailed, unpaired t-tests.

In agreement with our prior observation, the *formation* of filamentous cells predominantly depends on a minimal bacterial cluster size, independent of growth time or cluster distance (**Supplementary Figure S4A**), with ∼35 µm^2^ coverage in average. More strikingly, the formation of *interconnecting* filamentous cells also exhibits a strong cluster-size dependency, since we only observed those in much larger cell clusters (≥80 µm^2^). Upon discriminating between growth duration and spatial distances between cell clusters (**Figure 4B**), we observed that no *interconnecting* filamentous cells were formed after 6 h growth when separated by 14 µm, but otherwise the cluster sizes leading to either *forming* or *interconnecting* filamentous cells are comparable. Expectedly, the fraction of *interconnecting* cells increased significantly with growth duration (**Figure 4C**), mainly due to the associated increase in cell cluster size; after 18 h growth,∼50% of all filamentous cells were *interconnecting* cells connecting adjacent clusters. However, the fraction of *interconnecting* cells after 18h growth was significantly lower (∼16%) for clusters 14 µm apart compared to those separated by 9 µm. The dependency of cluster-*interconnecting* filamentous cells on cell cluster size and spatial distance supports our model that a diffusion-dependent process, such as DSF-dependent quorum sensing, triggers filamentous cell formation.

Particularly after 18 h growth, we observed many cell clusters interconnected with each other, suggesting that filamentous cells connect multiple adjacent clusters to form a large-scale network (**Figure 4D**). Upon analyzing the interconnections between all cell clusters (**Supplementary Figure S4B**), we were able to quantify the number of adjacent cluster interconnections per cell cluster (**Figure 4E**) for the two tested Au pattern distances (9 and 14 µm). The frequency distributions of the observed number of interconnections per cluster (**Figure 4E**) exhibited in case of 14 µm Au disk distances up to 6 interconnected cell clusters, whereas in case of 9 µm Au disk distances, the maximum observed number of interconnected clusters was 4. The difference in the degree of cluster interconnections as a function of cluster distances is more apparent upon comparing the average number of interconnected clusters (**Figure 4F**). Here, the average number of interconnections per cell cluster at the 14 µm cluster distance was ∼2-fold lower (∼0.8) than that observed for cell clusters spaced 9 µm apart (∼1.9). This striking result demonstrates that the degree of cell cluster interconnections is inverse to the spatial cluster distance, as we initially hypothesized. This result, together with the observation that multiple interconnections between large clusters were readily observable in biofilms (**Figure 4G**), support the idea that the formation of filamentous cells ultimately lead to large cluster networks that assemble into large-scale biofilms during its maturation.

## DISCUSSION

Our study elucidates the phenomenon by which cell filamentation occurs during bacterial biofilm formation, and reveals that their formation is crucial in the creation of large cell cluster networks that precede large-scale biofilms. Our developed platform to systematically investigate potential triggers and function of filamentous cell morphogenesis demonstrated that local cell abundance is the predominant parameter inducing cell filamentation, and that filamentous cell growth is directed towards proximate cell clusters in a cluster distance-dependent fashion, by which large interconnected cluster networks are formed. Both observed processes are consistent with the model of cell-cell communication in which a cell density-dependent gradient of diffusible signaling molecules drives morphogenesis *via* a quorum sensing mechanism.

### High affinity for cell adhesion to gold enables noninvasive, controlled spatial cell patterning

Selective and controlled spatial arrangement of cells and microcolonies are critical prerequisites for the investigation of dynamic biological phenomena of multicellular systems, such as cell plasticity, motility, morphogenesis, and cell-cell communication^35–37^. To this end, diverse approaches for selective cell organization with varying complexity and different surface chemical modifications have been previously developed^10,37–39^. To simplify the approach, we exploited the native ability of *X. fastidiosa* to bind to specific chemical moieties with high efficacy. Previous observations of the growth of *X. fastidiosa* on several different materials, such as Si, SiO_2_, InP, Au as well as various biotic surfaces mimicking the host environment, i.e., ethyl cellulose and acetate cellulose^4,31^, indicated a higher adhesion affinity to Au surfaces. Our results confirmed this feature, demonstrating that planktonic cells, and daughter cells after multiple cell divisions, predominantly adhered to gold. Our results further revealed a significant preference of cells to adhere to circular shapes over squared geometries, a characteristic that has also been previously observed for various biological interfaces^39,40^.

While bacterial cells predominantly adhered to gold deposits in our experiments, the number of cells adhering to the SiO_2_ interspace decreased with pattern distance. This rather non-intuitive observation might originate from the ability of *X. fastidiosa* to move along surfaces, at speeds up to 5 µm/min and against flow, *via* type IV pili-mediated twitching-motility^41^. Due to the taxis ability of *X. fastidiosa*, cells outside the Au areas might either randomly move until reaching the Au patterns, or even move directed to regions of higher cell amounts by chemotaxis (involving quorum sensing), which has been observed for other bacterial species^42–44^. The high affinity to gold is most probably mediated by its membrane protein methionine sulfoxide reductase that forms disulfide bonds to thiol groups on surfaces and adjacent cells^45,46^. In fact, membrane-associated thiol groups have been found in adhesion proteins and are key molecules involved in the adhesion mechanism of several bacterial species, including human pathogens^45,47,48^. Exploiting the strong interactions between gold and membrane-associated thiol groups^49^, coupled with the advantage that gold is biocompatible, chemically inert, and commonly used in biomedical applications^50,51^, our development provides a facile platform to create spatially well-defined cell adhesion patterns for noninvasive cell studies. Our methodology thus readily enables the systematic study of various complex phenomena involved in biofilm formation^52^ for a broad range of plant and animal pathogens^33,53,54^.

### Formation of cluster-interconnecting filamentous cells enables the creation of large biofilms

Our observed systematic dependency on cell cluster density and cluster distance in the formation of filamentous cells supports an earlier finding that first described the existence of filamentous cells and suggested that they might play a key role in the formation of large biofilm architectures by interconnecting cell clusters^11^.

The most intriguing result of our study revealed that filamentous cell growth is directed towards neighboring cell clusters to eventually integrate themselves into the other cluster. This growth behavior is unprecedented in bacterial pathogens, albeit suggested as *X. fastidiosa* cell clusters have been previously observed to be interconnected by filamentous cells^11^. Our observation that the filamentous cell growth is directed towards neighboring cell clusters depended strongly on the distance between the clusters, which in turn implies that the extent and direction of filamentous growth is governed by an extracellular regulatory mechanism. The possible linkage to quorum sensing in this process is attractive since DSF gradients produced by cell clusters could explain both the initiation of cell morphogenesis and the directional growth towards adjacent clusters.

Moreover, the potential triggering of cell filamentation by a quorum sensing mechanism might also be able to explain our observation that filamentous cell growth emerges preferentially from cells localized at cluster boundaries. In our experiments, only a small fraction of cells at cluster boundaries undergo cell morphogenesis, and resembles the result of our previous observation that similarly noted few filamentous cells in randomly nucleated bacterial clusters and small biofilms^11^. Taking into account that cell clusters become encapsulated in EPS during biofilm formation (i.e. loosely-bound EPS)^11,55^, cells at cluster boundaries might either not being entirely covered with EPS or the EPS layer is still thin enough to not fully act as DSF diffusion barrier, rendering the cells susceptible to regulatory quorum sensing (**Figure 5**). However, further investigation is warranted to elucidate if EPS can act as a potential DSF diffusion barrier.

**Figure 5:**
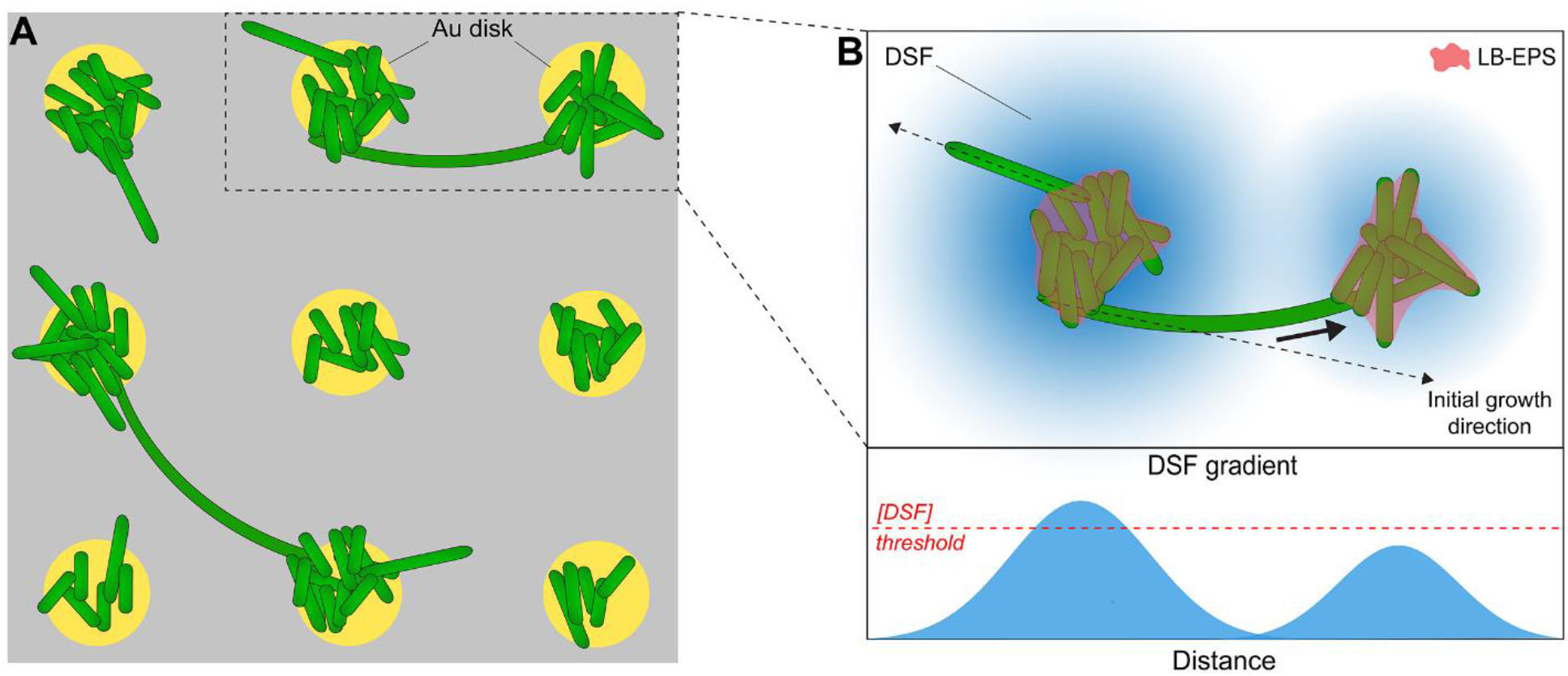
Proposed model of DSF-regulated cell morphogenesis and filamentous growth direction. (**A**) Schematic representation of experimentally observed filamentous cells interconnecting adjacent cell clusters. (**B**) High, cluster density-dependent local DSF concentration (in blue) triggers filamentous cell growth of non- or partly-EPS covered cells localized at cluster boundaries. While initial growth direction is random (dotted line arrows), filamentous cells encountering another DSF gradient of adjacent cell clusters continue growth and adjust growth direction (solid line arrow) towards the DSF gradient of adjacent cluster.

With respect to the underlying function of cluster-interconnecting filamentous cells, we observed that cells that had undergone such phenotypical morphogenesis eventually formed large networks of many interconnected bacterial clusters preceding the formation of larger biofilms. This finding confirms our original idea that cluster-interconnecting filamentous cells form a network that facilities spatial biofilm extension and its maturation^11^. During this process, continuous secretion of soluble EPS (i.e. S-EPS)^11^ by such filamentous cells would further provide an organochemical conditional film between the clusters that facilitates further cell adhesion^11^. The fact that that EPS-deficient mutants were not able to form biofilms support this model^56^.

While morphological plasticity and filamentous phenotypes have been reported for a variety of other bacteria, their morphogenesis was predominantly associated with adaptive responses to environmental changes and diverse forms of stress^6,16–19^. In contrast to this common belief, our results demonstrate that the formation of filamentous cells plays instead a vital role in *X. fastidiosa* biofilm formation *per se*. Analogous to our findings, filamentous cell formation, although in different contexts, has also been found essential for biofilm development and reinforcing cell surface adhesion for the pathogens *Vibrio cholera*^6^ and *Pseudomonas aeruginosa*^20^. As *X. fastidiosa* shares major genetic traits with other human and plant bacteria^14,15^, our findings might thus represent a conserved mechanism transferrable to other biofilm-forming bacterial pathogens.

### Bacterial density-dependent cell communication regulates filamentous cell growth

Biofilm-forming bacteria commonly communicate by means of a cell density-dependent mechanism known as quorum sensing^57–59^. Quorum sensing has been identified to regulate bacterial adhesion, virulence, gene expression, resistance, and other traits, rendering this process vital for the lifestyle adaptation in diverse pathogens^60–63^. In the case of *X. fastidiosa*, it has been similarly shown that cell-cell adhesion, virulence, and biofilm formation are regulated by the secretion of DSF molecules in a cell-density dependent fashion according to insect and plant hosts^21–23,25,34^.

The concept of cell density-dependent changes in cell behavior and phenotypes by DSF-based quorum sensing might underlie our observed cell density-dependent initiation of filamentous cell morphogenesis and directional cell growth^11,34,64^. In further support to this conjecture, previous observations in *X. fastidiosa* as well as in other bacteria and human cells have shown that quorum sensing mechanisms are activated upon exceeding a local DSF concentration threshold at regions of high cell densities^37,59,65–67^ Consistent with this model, a previous study using a *X. fastidiosa* Δ*rpfF* mutant blocked the production of DSF and exhibited significantly decreased cell adhesion to surfaces and was unable to form biofilms in an insect vector^22,23^. With respect to the overwhelming question of how many cells are required for cell density-dependent activation of cell morphogenesis, surprisingly, we observed a significant lower number of cells (∼15±5) compared to previous reported results^58,68,66^; for example, as many as ∼500 *P. aeruginosa* cells were necessary to detect a quorum sensing effect^68^.

Combining our observations with previous findings of quorum sensing processes in *X. fastidiosa* and other bacterial species, we propose a model of DSF concentration-dependent initiation of cell filamentation and cell cluster interconnection for *X. fastidiosa* (see **Figure 5**). Once a cell cluster is sufficiently dense to produce a local DSF concentration that passes a certain threshold, cells at the outer cluster boundaries, which are not or not fully encapsulated by EPS (i.e. LB-EPS^11^), undergo cell morphogenesis to filamentous cells. Our observation that a large number of filamentous cells stopped elongating in directions with either no adjacent cells or clusters consisting of a very few cells, implies that initial cell growth occurs in random directions. The fact that stark elongated filamentous, cluster-interconnecting cells often exhibited a curved shape with the cell pole oriented towards proximate clusters further suggests that filamentous cells can sense and respond to an increase of the surrounding DSF concentration by further growing towards the higher DSF gradient. The identified dependency on spatial proximity between bacterial cell clusters in quorum sensing efficacy has also been found earlier in other bacterial species^59,66,67^. Our observation that filamentous cells can change their growth direction towards adjacent clusters further suggests that the bacterial cell poles would likely contain the responsible quorum sensory proteins.

The question of which of the three currently known DSF molecules (XfDSF1, XfDSF2, CVC-DSF)^25^ might be responsible for triggering cell morphogenesis is difficult to answer, since filamentous cell growth is a rather new observation and the exact function of the DSF molecules within the *X. fastidiosa* lifecycle have not yet been fully identified. However, OMVs of all three *X. fastidiosa* strains (Temecula 1, 9a5C, Fb7) have been found to contain as cargo to neighboring and distant cells two of the hydrophobic DSF molecules (XfDSF2 and CVC-DSF), in addition to proteins for virulence and adhesion, such as lipases/esterases and adhesins, among others^25,34^. It has been suggested that the hydrophobic nature of *X. fastidiosa* DSF molecules allows them to embed within the cell membrane, and be subsequently distributed to other cells by release of OMVs^25,34^. Intriguingly, the OMVs secretion itself is also regulated by density-dependent, DSF-based communication^27^. OMVs might thus play a key role in initiating and driving filamentous cell growth by upon binding to the cell membrane of cells in clusters which are either not fully covered by EPS or residing outside the cluster boundaries. In this scenario, the DSF cargo, as well as adhesion-enforcing molecules, could be readily delivered to surrounding bacterial clusters. However, this model requires further systematic investigation in studies that involves altering DSF and/or OMV levels^21,56^.

Taken together, our study indirectly confirms the concept that DSF-based regulation is involved in the cell filamentation process during *X. fastidiosa* biofilm formation. This finding opens new directions that may lead to the identification of new targets in biofilm-forming pathogens and the design of quorum sensing inhibitors^33,69^ to inhibit biofilm formation^22,28,34^.

## METHODS

### Fabrication of Au micro array patterns

The Au micro patterns are fabricated by photolithography on cleaned borosilicate SiO_2_ substrates. Mask-free direct laser writing (DWL), equipped with a 380 nm solid state laser (Heidelberg Instruments µpg10; Power: 6 mW), was used on spin-coated AZ5214 photo resist (5,000 RPM for 50 sec, providing a resist layer with ∼1 µm thickness). After lithography, the patterns were developed using AZ351 (4:1) developer solutions for 15 s. Afterwards, 20 nm thick Au coating was deposited by electron beam physical vapor deposition (ULS600, Oerlikon Balzers, Liechtenstein) at 5 × 10^−7^ Torr. Finally, photo resist lift-off was carried by sonication with acetone for 2 minutes and rinsing with isopropanol and deionized (DI) water. The substrates were sterilized with oxygen plasma (SE80, Barrel Asher Plasma Technology, USA) for 10 min (100 mT, 50 sccm, 200 W) right before the bacterial adhesion experiments.

### Bacteria strains

*Xylella fastidiosa* pauca 11399 strain expressing soluble GFP was used in this study^11^. Periwinkle Wilt broth (PW) was used as bacterial growth media^70^. Bacteria extraction and pre-inoculum preparation was described elsewhere^11,32^.

### Bacterial growth

Bacterial inoculum with a concentration of 1 × 10^7^ CFU/mL from the pre-inoculum was used for the experiments as initial concentration for bacterial growth studies in PW broth media. The Au arrays were incubated for different growth times (specified in the respective manuscript text) in a bacterial stove (410/3NDR, Nova Ética, Brazil) at 28°C without culture media replacement. For Widefield microscopy (WFM) studies, the PW broth media was removed gently after certain growth times (6, 8, and 18 h). Samples were then washed twice with DI water to remove remaining chemical compounds of the culture media as well as non-attached bacteria. In a final step, the samples were dried gently with a nitrogen flow and temporarily stored at 4°C before fluorescence measurement.

### Wide-field epifluorescence microscopy

Dried bacteria samples of different growth times (6, 8, and 18 h) were measured using an epifluorescence microscope (Nikon TE2000U, USA) with a peltier-cooled back-illuminated EMCCD camera (IXON3, 1024 × 1024 pixels, Andor, Ireland) and a 60× water-immersion objective (CFI APO, NA 1.2, Nikon USA). GFP excitation and bacterial bright-field imaging was achieved by a 150 W Mercury-lamp with filter sets (AHF, Tübingen, Germany) for GFP (488 nm) and neutral density (ND8) filters, respectively. For each bacterial sample, a bright-field and a fluorescence image were taken sequentially. The images were merged and analyzed using Fiji/Imagej software.

### Confocal Laser Scanning Microscopy (CLSM)

For the *in-vitro* CLSM studies, the samples were placed inside a Teflon dish liquid cell (10 mm diameter and 5 mm in height), covered with a sterilized borosilicate cover glass. For each measurement, 400 µL of 4-times diluted *X. fastidiosa* 11399 inoculum was injected inside the liquid cell and incubated at 28°C for 14 h. CLSM measurements were performed using a Zeiss LSM780-NLO Confocal microscope (Carl Zeiss AG, Germany) with an 40x water-immersion objective (Plan-Apochromat, NA. 1.0, Zeiss) for *in-vitro* studies, and an 20× long-distance objective (Plan-Nanofluar, NA 0.5, Zeiss) for dried samples. The reflection of Au patterns and the fluorescence of GFP bacteria cells were simultaneously measured in two different channels. The GFP excitation was performed with a 488 nm laser line and the position of Au arrays were identified by the reflected laser. Imaging was performed with pinholes set to 1 airy unit for each channel, and with a 512 × 512 px image resolution. The images were merged and analyzed using Fiji/Imagej software.

### Image analysis

The measured fluorescence intensity, area of bacterial adhesion, as well as the integrated fluorescence density were extracted from raw fluorescence, reflective or bright-flied images using Fiji/Imagej. Background subtraction was performed on each individual fluorescence image.

## ACKNOWLEDGEMENTS

The authors would like to thank Prof. Steven E. Lindow (University of California, Berkeley) for fruitful discussions. The authors are greatly indebted to Vitor Pelegati (IFGW, UNICAMP) for his technical assistance during confocal microscopy measurements. This work was financially supported by the Brazilian agencies CNPq (441799/2016-7 and 429326/2018-1) and FAPESP (Grants number 2013/10957-0 and 2019/07616-3). S.A and A.M.S acknowledge FAPESP and CNPq, respectively, for funding their scholarships. The authors thank INFABIC/UNICAMP (FAPESP: 2014/50938-8, CNPq: 465699/2014-6), LAMULT, and the Device Research Laboratory (IFGW, UNICAMP) for granting access to their facilities.

## AUTHOR CONTRIBUTION

S.A., R.J. and M.A.C. conceived the project and designed the experiments. S.A., A.M.S., E.R.F.,M.S.S. and A.A.G.Z. optimized photolithography and Au coating conditions, prepared bacterial cultures and performed the experiments. H.F.C. and A.A.S. assisted with fluorescence microscopy and provided biological material. R.J. and M.A.C. directed the research. S.A., R.J. and M.A.C. analyzed the whole data and wrote the manuscript with input from all authors.

## Notes

### Competing Interest Statement

The authors have declared no competing interest.

