## Supplementary Figure for "Controlled spatial organization of bacterial clusters reveals cell filamentation is vital for *Xylella fastidiosa* biofilm formation"

### Supplementary Information

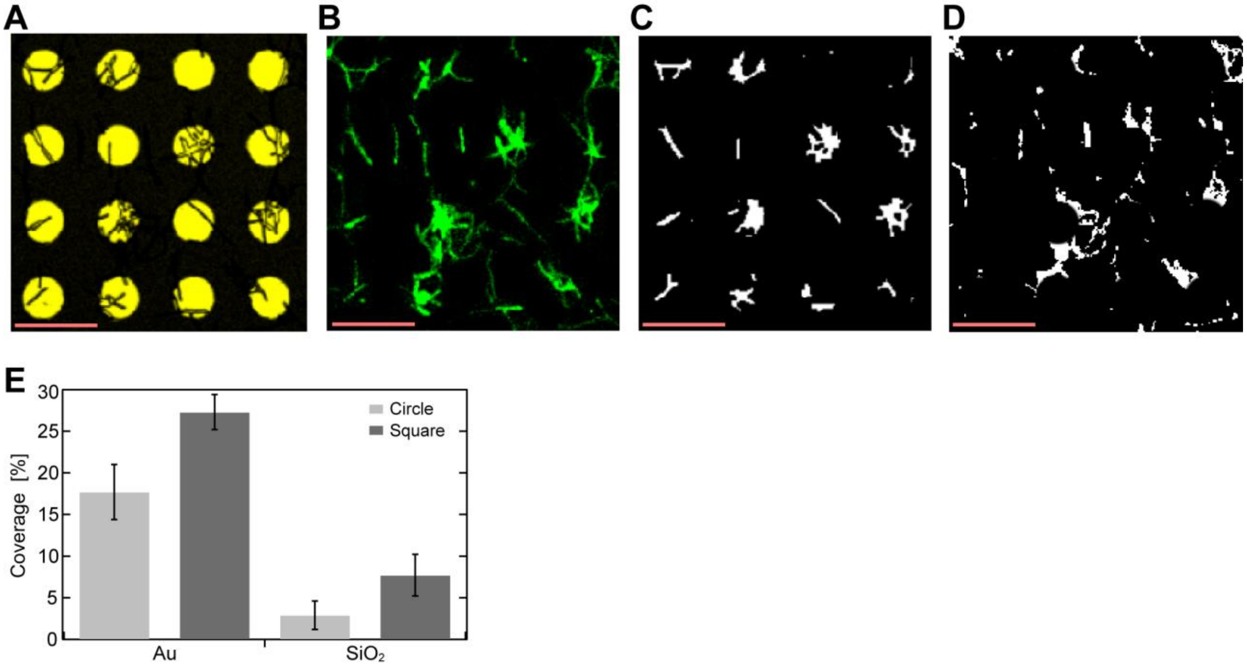


**Figure S1:** (**A**) Bright-field image of circular shaped Au patterns with adhered *X. fastidiosa*, and (**B**) corresponding fluorescence image after 18 h of growth. (**C**) Binary image of adhered *X. fastidiosa* on Au disks by extracting the areas of the Au disks shown in (A). (**D**) Binary image of adhered *X. fastidiosa* on $SiO_2$ surface by subtracting from the fluorescence image (B) the areas of the Au disks shown in (A). Scale bar depicts 20 µm. (**E**) *X. fastidiosa* coverage area (in % of total Au or $SiO_2$ area) on circular- and square-shaped Au surfaces, as well as on the $SiO_2$ surfaces, measured from data shown in Figure 1D data.

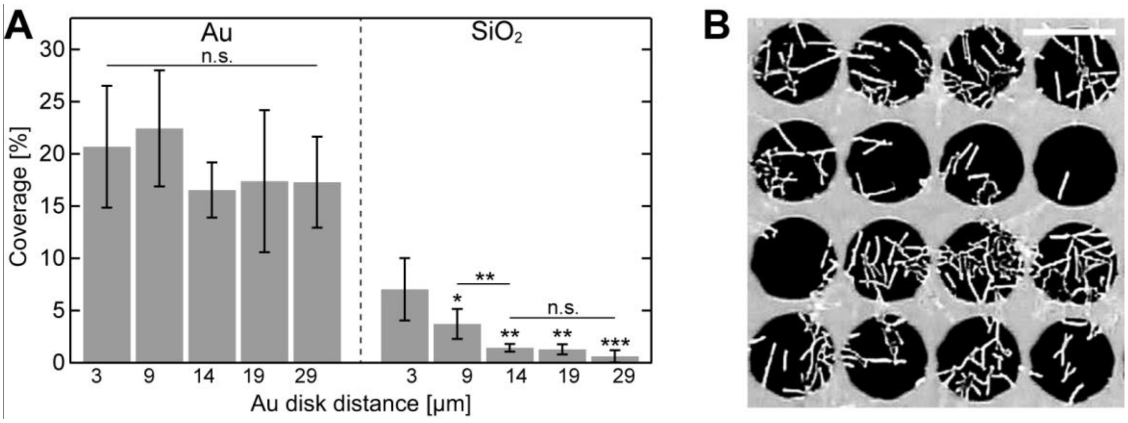


**Figure S2:** (**A**) *X. fastidiosa* coverage area (in % of total Au or $SiO_2$ area) on Au and $SiO_2$ surfaces, measured for different Au disk distances; related to Figure 2C. (**B**) Bright-field image of adhered *X. fastidiosa* on circular-shaped Au disks with 11 μm in diameter and 3 μm spacing between the disks after 8 hours of growth; scale bar depicts 15 μm.

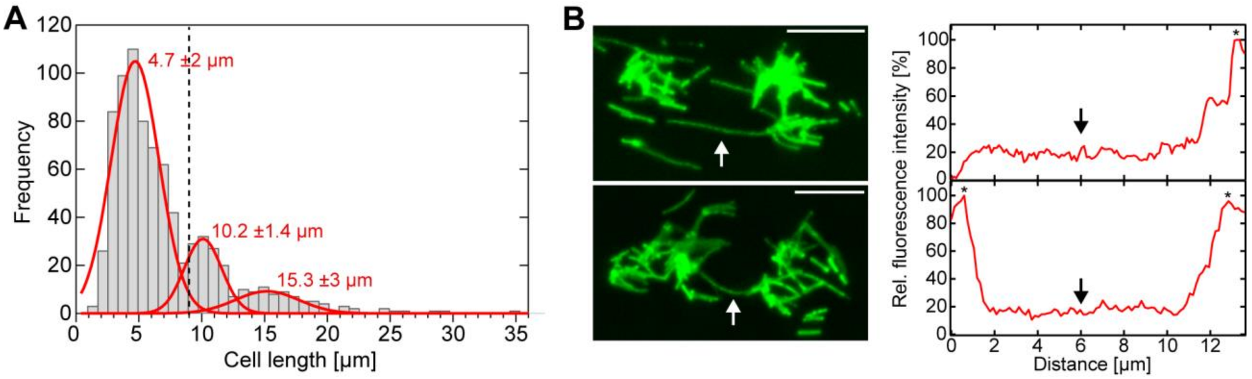


**Figure S3: (A)** *X. fastidiosa* cell length distribution (*N* = 833) pooled from experiments including both 6 and 18 h growth times, and 9 and 14 µm spacing between Au disks with 11 µm in diameter. Gaussian mixture model results in three distinct Gaussian fits (red lines), indicating three cell length populations. Dotted line indicates Gaussian peak -σ of the second population, determining a cell length threshold of 9 µm between typical cell lengths and cell length of filamentous cells. **(B)** Representative fluorescence images (left) of filamentous cells (indicated by white arrows) emanating from bacterial cell clusters, growing towards - and interconnecting - adjacent cell clusters. Corresponding background-corrected fluorescence intensity profiles (right) along the filamentous cells; scale bar depicts 10 µm. Asterisks on the intensity profiles represent the *X. fastidiosa* cluster boundary. Based on the results shown in (A) and (B), cell with lengths >9 µm and no significant GFP fluorescence signal variation along their cell body are considered as filamentous cells for analysis.

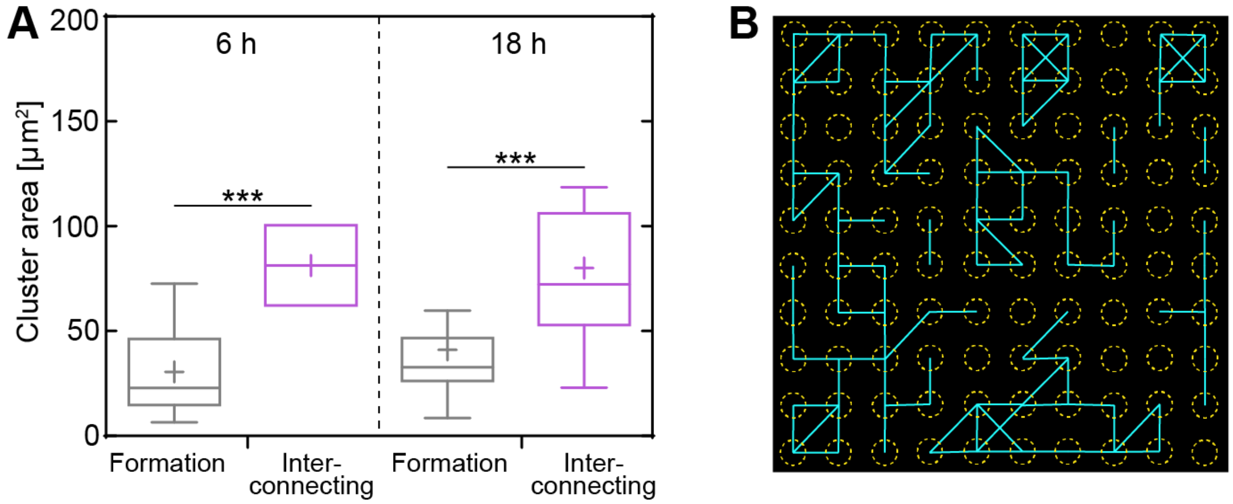


**Figure S4**: (**A**) Area distribution for bacterial cell clusters that solely form filamentous cells or form filamentous cell that interconnect neighboring clusters, for 6 h and 18 h growth time (Formation: N = 33 (6 h), N = 40 (18 h); interconnecting: N = 6 (6 h), N = 17 (18 h)). Statistical analyses consisted of unpaired, two-tailed t-tests (***$p < 0.001$). (**B**) Schematics of detected interconnections (cyan) of neighboring cell clusters (adhered to Au disks; yellow) by filamentous cells fluorescence image shown in Figure 4D.
